# Rehabilitation promotes the recovery of functional and structural features of healthy neuronal networks after stroke

**DOI:** 10.1101/141697

**Authors:** Anna Letizia Allegra Mascaro, Emilia Conti, Stefano Lai, Antonino Paolo Di Giovanna, Cristina Spalletti, Claudia Alia, Eros Quarta, Alessandro Panarese, Leonardo Sacconi, Silvestro Micera, Matteo Caleo, Francesco Saverio Pavone

**Affiliations:** Neuroscience Institute, National Research Council, Pisa, Italy; European Laboratory for Non-linear Spectroscopy University of Florence, Italy; Scuola Superiore Sant’Anna, Pisa, Italy; Scuola Normale Superiore, Pisa, Italy; National Institute of Optics, National Research Council, Italy; Ecole Polytechnique Federale de Lausanne (EPFL), Lausanne, Switzerland; Department of Physics and Astronomy, University of Florence, Italy

## Abstract

Rehabilitation is the most effective treatment for promoting the recovery of motor deficits after stroke. Despite its importance, the processes associated with rehabilitative intervention are poorly understood. One of the most challenging experimental goals is to unambiguously link specific circuit changes induced by rehabilitation to improved behavior. Here, we investigate which facets of cortical remodeling are induced by rehabilitation by combining optical imaging and manipulation tools in a mouse model of stroke. We demonstrate the progressive restoration of cortical motor maps and of cortical activity in parallel with the reinforcement of inter-hemispheric connectivity after rehabilitation. Furthermore, we reveal that the increase in vascular density goes along with the stabilization of peri-infarct neural circuitry at synaptic level. The present work provides the first evidences that rehabilitation is sufficient to promote the combined recovery of distinct structural and functional features distinctive of healthy neuronal networks.

## Introduction

Every year, several million stroke victims worldwide are impaired by long-term disability, imparting a large societal and personal burden in terms of lost productivity, lost independence and social withdrawal (Mozaffarian et al., 2015). Stroke survivors may suffer from lifelong losses in sensory, motor, cognitive, or emotional functioning, depending on the size and localization of the lesion. Persistent motor deficits generally occur after stroke damaged the motor-associated cortices. The recovery of motor function is often only partial, owing to the limited degree of spontaneous brain repair (see for example (Carmichael, Kathirvelu, Schweppe, & Nie, 2017)). On the other hand, strategies that promote brain plasticity, like pharmacological treatments and rehabilitation, can enhance neural rewiring and dramatically improve functional motor outcomes. Pharmacological treatment has been shown to aid the restoration of function in the peri-infarct area (for instance, see (Clarkson, Huang, Macisaac, Mody, & Carmichael, 2010)). A successful pharmacological approach is the modulation of contralesional hemisphere functionality, since disruptive influences of the intact hemisphere on the injured one (see for example (R. Y. Wang et al., 2012)) are often linked with worsened behavioral outcomes in stroke patients (Grefkes & Fink, 2011; Ward & Cohen, 2004). Another highly effective rehabilitative approach for restoring motor functions after stroke is physical training (Bütefisch, 2006; Jones & Adkins, 2015). Nevertheless, the lack of complete neuronal repair and functional recovery attained by pharmacological manipulation or motor training alone have led researchers to consider new combination approaches (Di Pino et al., 2014). The effectiveness of rehabilitative therapies can indeed be maximized through a combination of targeted physical exercise and other plasticizing treatments, such as cortical stimulation or pharmacological intervention (Adkins-Muir & Jones, 2003; Dancause & Nudo, 2011; Hesse et al., 2007; Plautz et al., 2003), aiming to specifically promote neuronal repair. A few animal studies provided useful insights into the anatomical adaptations in rodents and primates associated with combined therapies (Plautz et al., 2003; Wahl et al., 2014). For instance, axonal sprouting has been reported following sequential therapy with growth-promoting factors (like Nogo receptor antagonists (Fang et al., 2010; Lee, Kim, Sivula, & Strittmatter, 2004; Wahl et al., 2014) or chondroitinase ABC (Gherardini,Gennaro, & Pizzorusso, 2015; Soleman, Yip, Duricki, & Moon, 2012)) and intensive training. In a recent study on mice, Spalletti et al. (under revision) used a rehabilitation paradigm that combines motor training and pharmacological inhibition of the contralesional primary motor cortex (M1) with Botulinum Neurotoxin E (BoNT/E). BoNT/E is a bacterial enzyme that enters synaptic terminals close to the site of delivery and reversibly blocks neurotransmission by cleaving SNAP-25, a main component of the SNARE complex(Caleo et al., 2007). The injection of the toxin reduced the excessive transcallosal inhibition exerted from the healthy to the stroke side (Spalletti et al., under revision). The combination of motor training and silencing of the healthy hemisphere was superior to either treatment alone in promoting recovery of motor skills in stroke mice. Importantly, this combined therapy led to motor improvements that generalized to multiple motor tasks (Spalletti et al., under revision). The induction of a generalized functional gain, i.e. the recovery of motor functions beyond the ones that are trained, is crucial when evaluating the efficacy of rehabilitative therapies.

Based on this previous study, we herewith focused on the functional and structural plasticity triggered by combined rehabilitation. To overcome the limitation of *ex vivo* analysis, which can only provide a static view of brain dynamics (Misgeld & Kerschensteiner, 2006), we took advantage of *in vivo* optical imaging. In the last decade, fluorescence imaging provided valuable insights into spontaneous cortical plasticity after stroke. Murphy and colleagues performed a number of acute optical imaging studies showing the persistence of altered response selectivity of cortical areas over months after stroke (Brown, Aminoltejari, Erb, Winship, & Murphy, 2009; Harrison, Silasi, Boyd, & Murphy, 2013). In parallel, synaptic plasticity after stroke has been longitudinally evaluated by high resolution of two-photon fluorescence (TPF) microscopy (Craig E. Brown, Ping Li, Jamie D. Boyd, Kerry R. Delaney, & Timothy H. Murphy, 2007; Johnston, Denizet, Mostany, & Portera-Cailliau, 2013; Mostany et al., 2010; Sakadzic, Lee, Boas, & Ayata, 2015). *In vivo* TPF imaging has shown that the extent of synaptic recovery varies along the distance from the infarct core (Sigler & Murphy, 2010). Another *in vivo* TPF study found evidence for redistribution of blood vessel and dendrite orientation confined to the regions near the infarct border a few weeks after photothrombotic stroke (C. E. Brown, P. Li, J. D. Boyd, K. R. Delaney, & T. H. Murphy, 2007). Nevertheless, very little is known on the causative relationship between therapeutic intervention and brain plasticity after stroke. To this aim, the availability of a new generation of experimental tools (i.e. optogenetics) for manipulating neuronal activity provided new means of exploring (Ayling, Harrison, Boyd, Goroshkov, & Murphy, 2009; Lim, LeDue, Mohajerani, & Murphy, 2014) neuronal rewiring. In combination with the new generation of genetically encoded functional indicators, like GCaMP6 (Chen et al., 2013), optical imaging and optogenetics opened new opportunities to longitudinally dissect the underpinnings of therapy-aided motor recovery after stroke. To our knowledge, no investigation has yet described how rehabilitation affects cortical functional and structural plasticity *in vivo* and on the long-term scale. Many unanswered questions need to be addressed, like how rehabilitation after stroke longitudinally molds cortical functional maps, and how this is associated with modulation of inter-hemispheric connectivity. Furthermore, central issues like the impact of rehabilitation on neuronal structural plasticity are inadequately investigated, and yet never in vivo. Finally, it is still to be defined how neurons and vasculature act in concert to determine this complex and manifold reshaping.

In the present study, we dissected how rehabilitative treatment activates concurrent modalities of cortical plasticity in mice by using a multimodal approach that combines cutting-edge optical imaging and manipulation techniques. By longitudinal evaluation of cortical functional remodeling, we found that a combination of physical training with pharmacological manipulation of the contralesional activity is necessary to restore essential features of pre-stroke activation profiles, both in spatial and temporal terms. In the same animals presenting large-scale remapping of the injured hemisphere, we found a significant enhancement of transcallosal functional connectivity after one month of rehabilitative therapy. Furthermore, our experiments revealed an increase in the density of blood vessels primarily focused in the peri-infarct area, suggesting that the functional reshaping is supported by the neurovascular syncytium. Finally, as structural backbone of this modified functionality, we found in the same double-treated subjects that preservation of dendritic architecture goes along with stabilization of spines in the peri-infarct region. Overall, our study provides an unprecedented view of complimentary aspects of cortical anatomical and functional adaptation induced by rehabilitation after stroke.

## Results

### Experimental design

We used a photothrombotic model of focal stroke applied to the right M1 of GCaMP6f or GFPM mice (STROKE, ROBOT and REHAB groups). The combined rehabilitation paradigm of the REHAB group consists in a combination of highly repeatable motor training of the mouse forelimb in a robotic platform (M-Platform (C. Spalletti et al., 2014)) and reversible pharmacological inactivation of the healthy, contralesional hemisphere via the synaptic blocker Botulinum Neurotoxin E (BoNT/E; see Supplementary Fig. 1A). The ROBOT group performed the same daily training on the M-platform as REHAB mice, but without pharmacological treatment. Rehabilitation-associated training on the M-Platform consisted of repeated cycles of passively actuated contralesional forelimb extension followed by its active retraction triggered by an acoustic cue (Fig. 1A and Supplementary Fig. 1B).

ROBOT and REHAB mice were trained for 4 weeks starting 5 days after injury (Supplementary Fig. 1C), in line with the overall consensus that the initiation of rehabilitative training 5 or more days after stroke is mostly beneficial and has no adverse effects(Krakauer, Carmichael, Corbett, & Wittenberg, 2012). The motor task was rapidly learned and easily performed. Indeed, the performances, evaluated by the amplitude and slope of the force peaks exerted during the voluntary forelimb-pulling task, were not significantly different across groups, neither within a week nor across weeks of training (Supplementary Fig. 1D). The treatment induced a generalized improvement in motor performance, as assessed by the Schallert Cylinder test(Schallert, Fleming, Leasure, Tillerson, & Bland, 2000) one month after the lesion. While the STROKE mice showed a significant asymmetry in forelimb use, this impairment was no longer present in REHAB mice at 30 days post-stroke (Supplementary Fig. 1E).

### Combined rehabilitation treatment restores cortical activity patterns disrupted by stroke

To assess the extent to which the improved motor performance was accompanied by large- and small-scale cortical plasticity, we investigated the remapping of motor representations, vascular network remodeling and synaptic dynamics. First, we evaluated how rehabilitation modulated cortical functionality over the meso-scale. We implemented an integrated system for simultaneous imaging of the calcium indicator GCaMP6f and recording of forces applied by the contralesional forelimb during the training sessions on the M-Platform (Fig. 1A-C). Calcium imaging was used as a measure of the cortical activity in the brains of GCaMP6f mice. Wide-field calcium imaging showed that a small area located in the motor-sensory region reproducibly lit up in CTRL mice during the M-platform forelimb retraction movement (an example of cortical activation is reported in Fig. 1D and Supplementary Fig. 2A). On the contrary, a large area covering most of the cortical surface of the injured hemisphere was activated synchronously in non-treated (STROKE) mice one month after stroke while performing the task. We tested if 4 weeks of motor training on the M-platform (ROBOT group) could restore pre-stroke conditions. We observed that motor rehabilitation without pharmacological intervention resulted in a diffuse activation, originating from the peri-infarct cortex (Supplementary Fig. 2A). Remarkably, in comparison to STROKE and ROBOT mice, REHAB mice showed a more spatially restricted zone of activation that was similar to the pattern seen in healthy controls (CTRL) in terms of location, timing and amplitude (Fig. 1D-H, Supplementary Fig. 2A-D).

**Figure 1:**
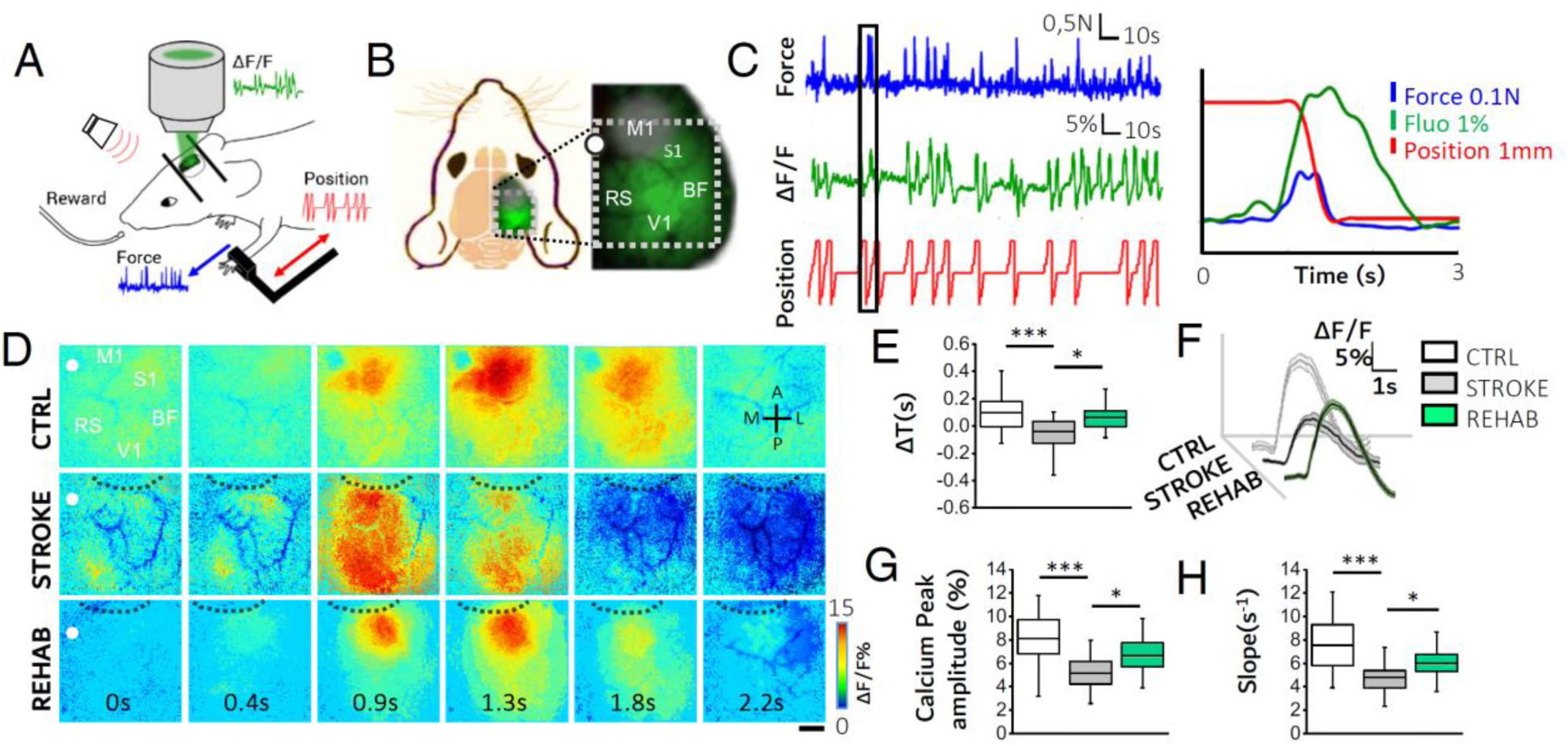
Combined rehabilitation counteracts cortical dedifferentiation and restores cortical activation profiles in the post stroke, peri-infarct region. (A) A schematic representation of the M-Platform used for rehabilitative training and simultaneous calcium imaging. Examples of force (blue), position (red) and ΔF/F traces (green) are reported. (B) A schematic representation of the field of view (i.e. the area within the dotted white square) used for wide-field calcium imaging in GCaMP6f mice. The gray circle on the M1 region highlights the location and approximate extent of the lesion. M1, primary motor area; S1, primary somatosensory area; BF, barrel field; V1, primary visual cortex; RS, retrosplenial cortex. The white dot indicates the bregma. (C) An example of the force trace (blue trace), the simultaneously recorded calcium fluorescence trace (green trace) and the position of the handle (red trace). The graph on the right shows an overlap of the simultaneously recorded traces that correspond to the black box on the left. (D) An image sequence of cortical activation as assessed by calcium imaging during pulling of the handle by the contralateral forelimb, from 0.4 s before to 1.8 s after the onset of the force peak. Each row shows a representative sequence from a single animal of each group. Black dashed lines define the lesion borders. Scale bar, 1mm. (E) The delays in cortical activation in caudal regions in response to contralateral forelimb retraction are shown for the 3 groups (Nmice_CTRL_= 4, Nmice_STROKE_ = 6, Nmice_REHAB_= 6; ΔT_CTRL_ = 0.10 ± 0.03 s, ΔT_STROKE_ = −0.04 ± 0.02 s, ΔT_REHAB_= 0.06 ± 0.02 s; Kruskal-Wallis One Way followed by Tukey′s test: *** *P* = 0.006, * *P* = 0.016). (F) Average calcium traces recorded in the region of maximal activation, ROIg (see Methods), during contralateral forelimb retraction for the three experimental groups; the shadows indicate the SEM values. (G) The graph shows the maximum of the fluorescence peaks from the same calcium traces as in (F) (Peak amplitude_cTRL_ = 8.1 ± 0.5 %; Peak amplitude_STROKE_ = 5.2 ± 0.3%; Peak amplitude_REHAB_ = 6.6 ± 0.3%; one-way ANOVA followed by Tukey′s test: *** *P* = 0.000005, * *P* =0.017). (H) The graph shows the slope (average ± SEM) of the fluorescence in the rising phase of the trace (Slope_CTRL_ = 7.5 ± 0.5 s^−1^; Slope_STROKE_ = 4.8 ± 0.2 s^−1^, Slope_REHAB_ = 6.0 ± 0.2 s^−1^; Kruskal-Wallis One Way followed by Tukey′s test: *\*\*\* P* = 0.00004, * *P* = 0.015).

By overlapping the forelimb retraction-triggered cortical maps obtained on every day of the training week, we quantified the location and origin of activation of the calcium waves (Supplementary Fig. 2B-D). Stroke shifted and expanded the location of the retraction task motor representation from M1 toward more caudal brain regions not specifically associated with motor control in STROKE mice (Supplementary Fig. 2C, 2^nd^ panel). ROBOT mice showed a cortical activation that positively correlated with proximity to the stroke lesion, but more diffuse compared to CTRL mice (Supplementary Fig. 2C, 3^rd^ and 1^st^ panel, respectively, and Supplementary Fig. 2D). Interestingly, the spread of cortical activation in REHAB mice was further reduced towards pre-stroke conditions (Supplementary Fig. 2C 4^th^ panel and Supplementary Fig. 2D). In most cases (5 out of 6 REHAB mice), the improvement in cortical activation patterns was associated with increased functional connectivity between the primary and secondary motor output areas at the end of the training period (REHAB/4W) compared to the beginning (REHAB/1W), towards healthy (CTRL) conditions (Supplementary Fig. 2E). Our results indicate that the combined rehabilitative treatment focalizes the cortical activation to the peri-infarct region while limiting the concurrent recruitment of distant locations.

We hypothesized that the spatial spread of calcium waves implied a synchronicity in the activation of the rostral peri-infarct network and the caudal inter-connected region. By analyzing the temporal profile of forelimb retraction-triggered cortical activation, we found that the delay between the maximum calcium peak of the rostral and caudal regions was reversed in STROKE versus CTRL mice, indicating that the activation of the caudal region somewhat preceded the activation of the peri-infarct areas in STROKE mice (Fig. 1E). Longitudinal imaging showed that the temporal dynamics of calcium activation was gradually recovered during the 4 weeks of training in REHAB mice (Supplementary Fig. 2F, left panel).

The spatio-temporal profile of the REHAB group closely resembled CTRL animals, suggesting that the motor output pattern underlying the forelimb retraction primarily involved the peri-infarct region. Furthermore, we found that the combined treatment profoundly affected the neuronal activation patterns of REHAB mice. Indeed, the average calcium traces in Fig. 1F show the differences in the calcium dynamic measured in the region of maximum activation (*ROIg*, see Supplementary Fig. 2B) during contralateral forelimb retraction in CTRL, STROKE, and REHAB mice.

While a significant reduction in the amplitude of cortical activity during forelimb retraction was evident one month after stroke in the STROKE group, REHAB mice partially recovered to pre-stroke conditions (Fig. 1G). The combined treatment led to a progressive increase in the amplitude of the calcium response over the training period (Supplementary Fig. 2F, middle panel).

Moreover, REHAB animals showed progressively steeper slope of calcium activation during the rehabilitation period (Supplementary Fig. 2F, right panel). As compared to STROKE animals, at the end of the training period REHAB mice demonstrated a faster calcium activation curve (Fig. 1H).

We observed that the training on the M-Platform for 4 weeks alone (ROBOT mice) resulted in a small but not significant improvement in calcium kinetics parameters compared to STROKE mice (Supplementary Fig. 3).

Conversely, the combined treatment (REHAB mice) promoted a solid rescue of calcium kinetics significantly different from STROKE mice towards heathy conditions (Fig. 1E-H). Based on the modest changes induced on cortical functionality in the ROBOT group, the following experiments were focused exclusively on the combined treatment (REHAB group).

In brief, our results show that combined rehabilitative treatment is necessary to promote the recovery of the temporal and spatial features of cortical activation patterns in the peri-infarct region that are associated with the execution of a motor task.

### Inter-hemispheric functional connectivity is improved by combined rehabilitation

Having analyzed the ipsilateral adaptation of cortical activity, we next evaluated how distal interconnected regions, such as the contralesional motor cortex, were affected by our therapeutic intervention. Several studies in mice and humans have shown that interhemispheric M1 connectivity is reduced after stroke (Bauer et al., 2014; van Meer, van der Marel, Otte, Berkelbach van der Sprenkel, & Dijkhuizen, 2010; van Meer, van der Marel, Wang, et al., 2010). Increased interhemispheric connectivity is hypothesized to positively correlate with the recovery of motor performance in the subacute stage after stroke in humans (Carter et al., 2010). Based on this hypothesis, we tested whether transcallosal connectivity between homotopic areas was modulated by the double rehabilitative treatment by using an all-optical approach that combined optogenetic activation of the intact M1 with calcium imaging of the cortex in the injured hemisphere. In these experiments, the spared, left M1s of GCaMP6f mice were injected with AAV9-CaMKII-ChR2-mCherry to induce the expression of ChR2 in excitatory neurons (Fig. 2A).

One month after stroke, we optogenetically stimulated the intact M1. To avoid ChR2 stimulation while exciting GCaMP6f fluorescence, we partially occluded the 505 nm LED in our custom-made wide-field microscope (Fig. 2B-C). Optogenetic stimulation was achieved with a second excitation path in which a 470 nm laser was focused on the AAV-transfected region via acousto-optic deflectors (AODs).

**Figure 2.**
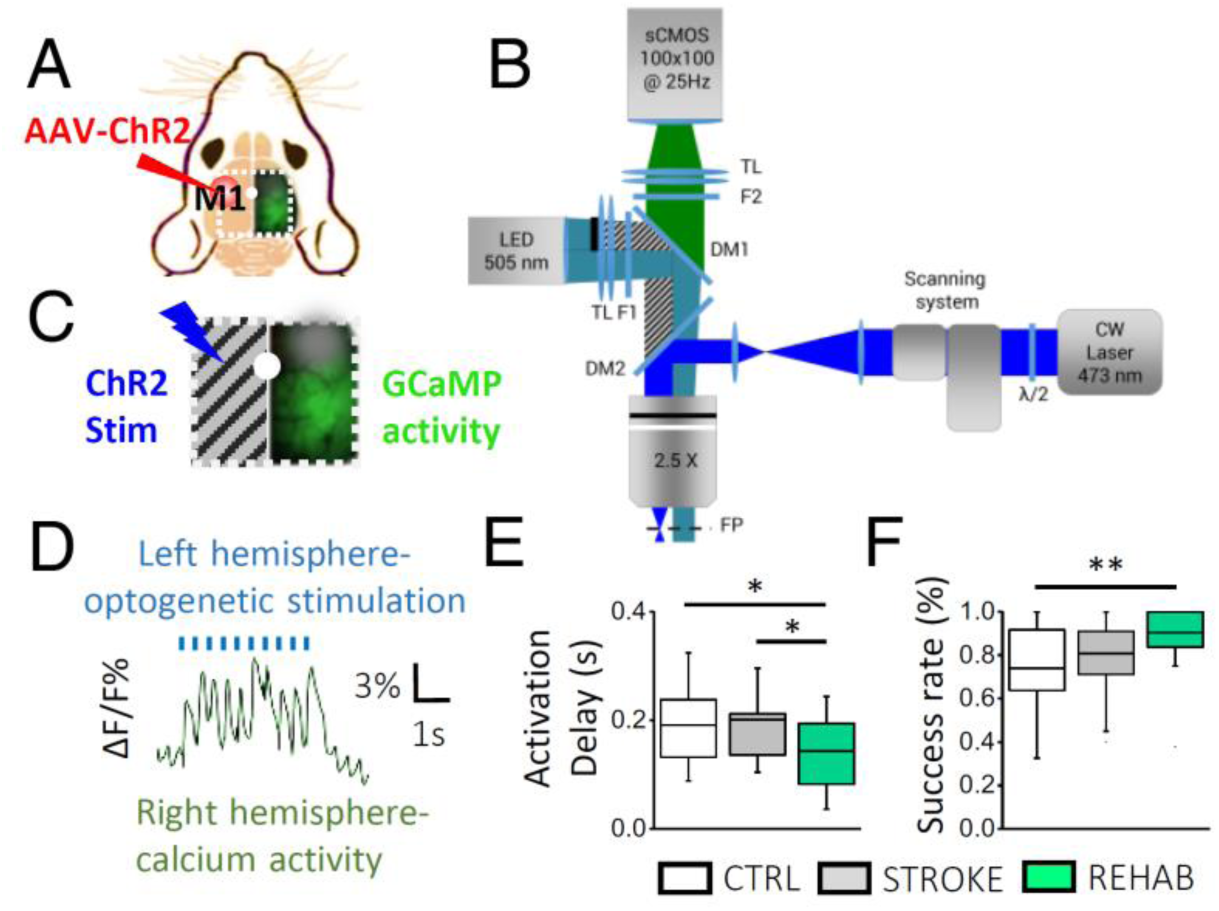
Combined rehabilitation strengthens interhemispheric connectivity after stroke. (A) A schematic representation of the field of view (i.e. the area within the white dotted square) for the all-optical investigation of inter-hemispheric functional connectivity in GCaMP6f mice. The red circle in the left hemisphere indicates M1 injected with AAV-ChR2-mCherry; the gray circular “cloud” labels the site of the stroke lesion in the right hemisphere. The white dot indicates bregma. (B) A schematic representation of the wide-field microscope with a double illumination path that allows simultaneous light stimulation for optogenetics and fluorescent recording for calcium imaging. The hemi-occlusion of the 505nm LED, displayed by the black line on the upper half of the LED illumination path, avoids spurious ChR2 stimulation during calcium imaging. TL, tube lens; F1/2, band-pass filters; DM1/2, dichroic mirror; FP, focal plane. (C) The panel shows the simultaneous ChR2 laser stimulation on the left hemisphere (not illuminated by LED light) and the detection of the evoked GCaMP6f fluoresc ence on the right side. (D) A trace of optogenetically elicited calcium activity in the right cortex. (E) Box and whiskers plot showing the delay between the onset of left hemisphere laser stimulation and the peak of the calcium responses in the right hemisphere (average ± SEM; Nmice_CTRL_ = 4, Nmice_STROKE_ = 3, Nmice_REHAB_ = 3; Activation Delay_CTRL_ = 0.19 ± 0.02 s, Activation Delay_STROKE_ = 0.20 ± 0.02 s, Activation Delay_REHAB_ = 0.14 ± 0.01 s; one way ANOVA post hoc Fisher, *P* <0.05 for all comparisons;). (F) The success rate (SR) of laser stimulation, calculated as the number of times the laser stimulation successfully triggered contralateral activation over the total number of stimulation trials (average ± SEM, same Nmice as in (E); SR_CTRL_= 0.74 ± 0.04, SR_STROKE_= 0.81 ± 0.03, SR_REHAB_= 0.90± 0.03; ANOVA with post hoc Bonferroni test: *P* < 0.01).

Laser stimulation of the contralesional M1 reproducibly triggered calcium waves in the injured hemisphere (Fig. 2D, Supplementary Video 1), beginning mainly in the homotopic M1 of CTRL mice, or in the peri-infarct area of STROKE and REHAB animals (Supplementary Fig. 4).

Calcium waves then propagated to functionally connected regions that were located either anterior or posterior to the primary source of activation in all 3 groups of animals (Supplementary Video 1). By visual inspection, the temporal dynamics of the calcium waves were comparable between CTRL and STROKE mice (Supplementary Fig. 4).

Indeed, by quantifying the delay between the start of the optogenetic stimulation on the left hemisphere and the peak of calcium activity in the right hemisphere, we found no significant difference between the STROKE and CTRL mice (Fig. 2E). In contrast, this delay was significantly reduced in REHAB mice as compared to both CTRL and STROKE mice (Fig. 2E). In addition, the success rate of contralateral activity in response to optogenetic stimulation was higher in REHAB mice than in CTRL and STROKE animals (Fig. 2F). Therefore, this series of all-optical experiments suggests that combined rehabilitation strengthens the functional connectivity between the spared motor cortex and the perilesional cortex one month after stroke.

### Combined rehabilitative treatment does not recover vessel orientation but stimulates angiogenesis in the peri-infarct area

We next assessed whether vascular reorganization structurally correlated with of the functional remapping that occurred at the local and inter-hemispheric levels in REHAB animals after stroke. To test this, we longitudinally observed mouse cortices of all groups under both the cranial window and the thinned skull preparations in the month following the induced stroke, and found that the injured area consistently shrunk owing to the collapse of dead tissue in STROKE and REHAB mice (Fig. 3A and 4A). In addition, a very bright area appeared in the peri-infarct region of both groups of GCaMP6 mice, possibly representing an excitotoxic response associated with calcium dysregulation (Ankarcrona et al., 1995) elicited by the photothrombotic stroke (see the left panel of Fig. 3A; 13 out of 14 STROKE and REHAB mice). The enhanced brightness gradually diminished and disappeared after the acute period (6-19 days after injury in STROKE and REHAB mice; see example in Fig. 3A, right panel), and was accompanied by a large shrinkage of the necrotic tissue. The consequent displacement of the peri-infarct area was associated with a substantial remodeling of blood vessels (white arrowheads in Fig 3A), in agreement with previous studies (C. E. Brown et al., 2007).

To quantify the structural reshaping associated with stroke and rehabilitation, we performed 3D reconstructions of the cortical vasculature. Specifically, after the last motor training session, we stained the brain vasculature of GFPM mice with a vessel-filling fluorescent dye, albumin-fluorescein isothiocyanate (FITC, Fig. 3B). By performing TPF microscopy on Thiodiethanol (TDE) cleared cortical slices (see Methods), we obtained high-resolution 3D maps of blood vessels of the injured, right hemisphere (Fig. 3C, Supplementary Video 2). A strong orientation gradient of blood vessels induced by the collapse of dead tissue in the ischemic core was evident in both the STROKE and REHAB groups. Indeed, combined rehabilitative treatment did not rescue random vascular orientation of pre-stroke conditions in regions proximal (<500 μm) or distal (1000 to 1500 μm from core) to the stroke core in REHAB mice (Fig. 3D). On the other hand, the density of blood vessels near the infarct site was significantly higher in REHAB mice than in CTRL and STROKE groups (Fig. 3E, left panel). We further evaluated the vascular density over the entire cortical depth to define each layer-specific contribution to the overall increase in blood vessel density in REHAB animals. The blood vessel distribution was uniform in both CTRL and STROKE mice. Increases in density were localized mainly to the middle and deeper layers of the cortex (300-700 μm deep) in the REHAB group(Fig. 3E, middle panel). We found that this layer-specific increased vascular density in the REHAB animals, which was possibly due to treatment-induced angiogenesis, was mainly observed in the region proximal to the stroke core (Fig. 3E, right panel). Finally, we found no significant differences in the distribution of lumen sizes of cortical blood vessels for the three experimental groups (Fig. 3F), ruling out possible influences of vasodilation or vasoconstriction in the evaluation of the density.

**Figure 3.**
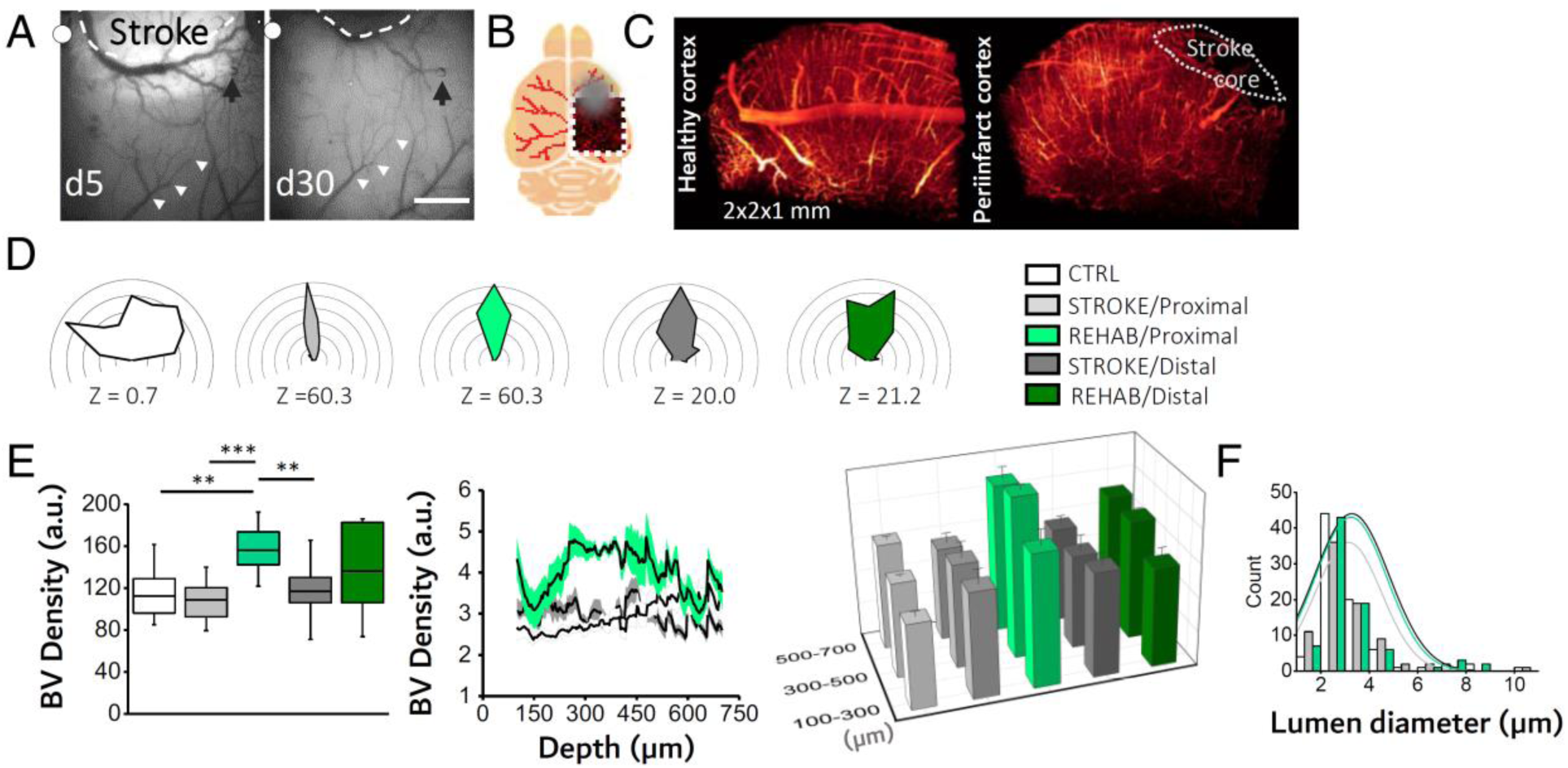
Combined rehabilitation promotes angiogenesis in the peri-infarct area. (A) Brightfield images showing the cortical vasculature following a stroke under a thinned skull preparation. The dotted line emphasizes the profile of a large blood vessel shifting toward the stroke core from 5 days to 30 days after stroke; a similar shift is highlighted by the white arrowheads on a blood vessel distal to the injury site. The black arrow points to an internal reference on the image. The white dots indicate bregma. (B) A schematic representation of the fixed brain of a GFPM mouse, in which the vasculature is labeled with TRITC (red lines). The dotted square highlights the imaged area; th e gray circle labels the stroke location. (C) A 3D meso-scale reconstruction of the vasculature in the healthy contralesional cortex (left) and in the peri-infarct area (right) from TDE-cleared cortical slices obtained from a STROKE GFPM mouse, one month after injury. The white dotted region highlights the absence of blood vessels in the infarct core. (D) The polar plot shows the distribution of blood vessel orientation in regions proximal and distal to the core for each experimental group measured after the last motor training session (Nmice_CTRL_ = 3; Nmice_STROKE_ = 3; Nmice_REHAB_ = 3). The plots are oriented as in Fig. 3A; the lesioned area is toward the uppermost centre of the plot. Z scores, calculated by the Rayleigh Test for circular statistics, are reported for each experimental class below the plot. (E) Blood vessel (BV) density analysis. (*Left*) The histogram shows the blood vessel density (average ± SEM) in proximal and distal regions from the stroke core in each experimental group ( BV Density_CTRL_ = 11.2 ± 0.7 a.u.; BV Density_STROKE Proximal_ = 10.9 ± 0.6; BV Density_STROKE Distal_ = 11.7 ± 0.6; BV Density_REHAB Proximal_ = 15.6 ± 0.7; BV Density_REHAB Distal_ = 13.6 ± 1.1; one-way ANOVA with post hoc Bonferroni test: *** *P* <0.001, ** *P* <0.01, * *P* <0.01). (*Middle*) The traces indicate the blood vessel density with respect to brain cortex depth in each experimental group. Gray shadows represent the SEM. (*Right*) A 3D graph comparing the average blood vessel density grouped by cortical depth (100-300 μm from stroke core: BV Density_CTRL_ = 2.5 ± 0.2; BV Density_STROKE Proximal_ = 3.1 ± 0.4; BV Density_REHAB Proximal_ = 3.9 ± 0.5; BV DensityS_TROKE Distal_ = 3.1 ± 0.4; BV Density_REHAB Distal_ = 2.9 ± 0.4; 300-500 μm from stroke core: BV Density_CTRL_ = 2.8 ± 0.1; BV Density_STROKE Proximal_ = 3.1 ± 0.5; BV Density_REHAB Proximal_ = 4.7 ± 0.2; BV Density_STROKE Distal_ = 2.8 ± 0.5; BV Density_REHAB Distal_ = 3.5 ± 0.2; 500-700 μm from stroke core: BV Density_CTRL_ = 3.1 ± 0.1; BV Density_STROKE Proximal_ = 2.8 ± 0.3; BV Density_REHAB Proximal_ = 4.4 ± 0.4; BV Density_STROKE Distal_ = 2.8 ± 0.3; BV Density_REHAB Distal_ = 3.6 ± 0.1; 300-500 μm from the stroke core: *P* <0.05 for REHAB Proximal vs STROKE Proximal, STROKE Distal, and CTRL; 500-700 μm from the stroke core: *P* <0.05 for REHAB Proximal vs STROKE Proximal and STROKE Distal, one-way ANOVA with post hoc Bonferroni test). (F) The histogram shows the distribution of lumen diameter of blood vessels for CTRL mice and in the proximal region of STROKE and REHAB mice.

In conclusion, by combining *in vivo* and *ex vivo* imaging methods, we generated 3D cortical vascular maps and observed a pro-angiogenic effect of the combined rehabilitative treatment on the vasculature of the peri-infarct area.

### Combined rehabilitation preserves pyramidal neurons architecture and promotes the stabilization of synaptic contacts in the peri-infarct area

In addition to vascular reshaping, we explored finer neuro-anatomical adaptations to define their contribution to changes in cortical activity after stroke. Heightened spontaneous structural remodeling has been observed in animals in both presynaptic (axonal fibers and terminals) and postsynaptic (dendrites and dendritic spines) neuronal structures in the peri-infarct cortex in the weeks following stroke (Brown, Boyd, & Murphy, 2010; Carmichael, Wei, Rovainen, & Woolsey, 2001; Dancause et al., 2005; Hsu & Jones, 2006; Ueno et al., 2012). In the next set of *in vivo* experiments, we aimed at defining if rehabilitation after stroke was associated with an alteration in synaptic-level cortical circuits, and how this change was modulated by distance from the core. We performed TPF microscopy of pyramidal apical dendrites and axons (up to 100 μ deep relative to the pial surface) at increasing distances (up to 4 mm) from the stroke core in the cortex of GFPM mice. The imaging area overlapped with the region visualized in GCaMP6f mice (Supplementary Fig. 5A). The orientation of dendrites and axons was evaluated on the 4^th^ week after injury in STROKE and REHAB mice.

In the same animals studied on the vascular imaging experiments, the reorganization of cortical tissue (Fig. 4A, left panels) that led to a considerable alignment of blood vessels towards the stroke core produced an analogous re-orientation of cortical dendrites and axons in the peri-infarct area of STROKE mice (white arrow in Fig. 4A, right panels). Reorientation was less pronounced at distal regions from the ischemic core (>1 mm; Supplementary Fig. 5C). In contrast, the change in organization of pyramidal processes was not visible in REHAB animals (Fig. 4B and Supplementary Fig. 5B) even though stroke profoundly altered the spatial distribution of blood vessels around the stroke core in the same mice. Indeed, the randomness in dendrite orientation typical of healthy subjects was preserved in REHAB mice as compared to the STROKE group (Fig. 4B).

**Figure 4.**
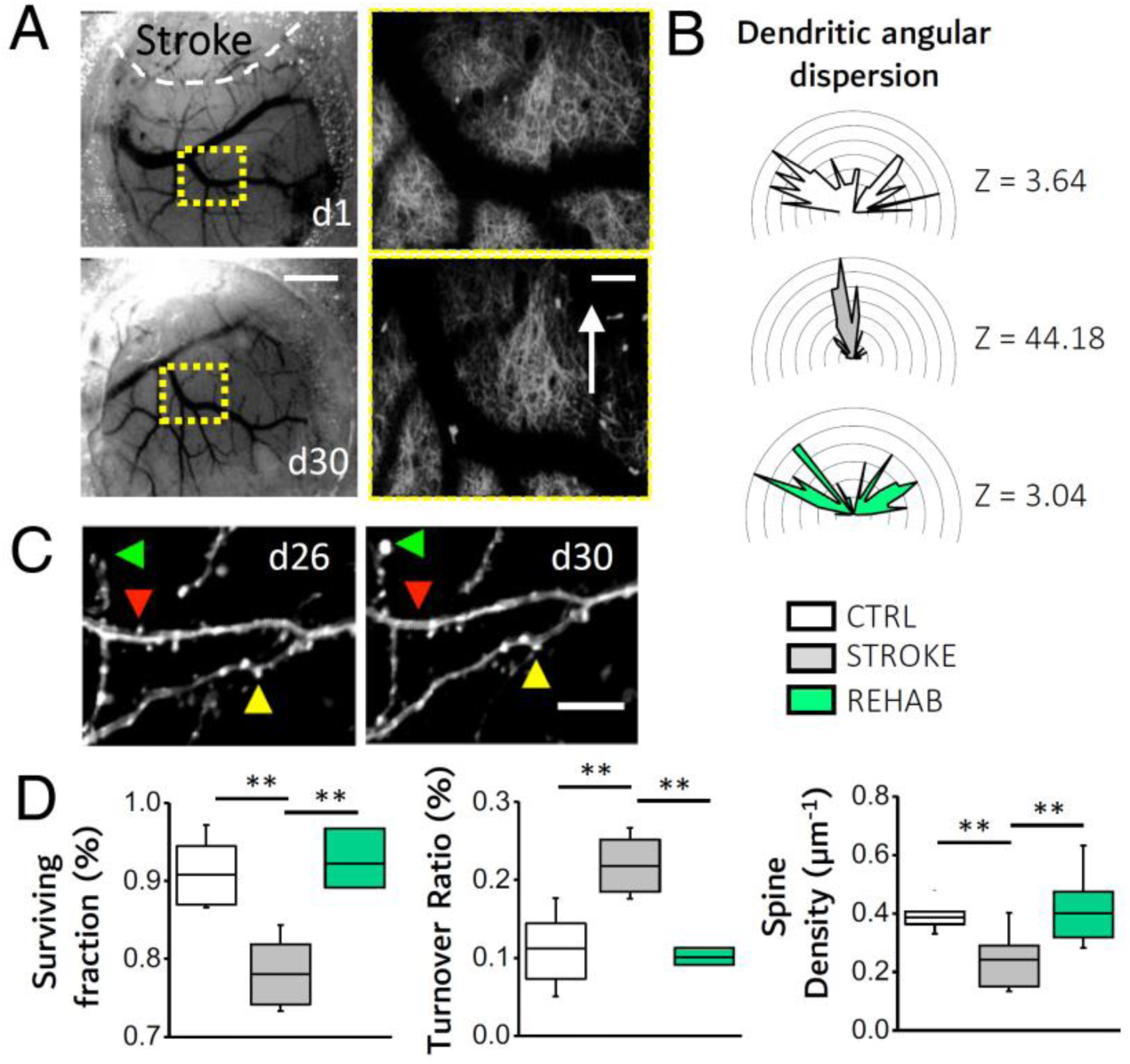
Combined rehabilitation affects dendritic orientation and stabilizes spine turnover. (A) Brightfield images showing the cranial window 1 day and 30 days after stroke (STROKE group). The rostral shift of cortical tissue due to the shrinkage of the stroke core is highlighted by the displacement of a reference point (i.e. blood vessel branching point) framed by the dotted yellow square. Scale bar = 1 mm. The panels on the right show the stitched two-photon images (4×4 Maximum Intensity Projections, MIPs of 130×130×50 μm^3^) acquired within the region framed by the respective yellow squares in the left panels. The white arrow highlights the alignment of dendrites and axons towards the stroke core. Scale bar = 100 μm. (B) Polar plots showing the angular distribution of dendrites in the peri-infarct area. The plots are oriented as in Fig. 4A, where the lesioned area is toward the uppermost center of the plot (for all panels, Nmice_cTRL_ = 5, Nmice_STROKE_ = 4, Nmice_REHAB_ = 3). Z scores, calculated by the Rayleigh Test for circular statistics, are reported for each experimental class on the right of the polar plot. (C) MIPs of two-photon stacks (z depth 8 μm) of dendritic branches at 26 and 30 days after stroke (STROKE group). The arrows point to a newly formed (green), lost (red), and stable (yellow) spine. Scale bar = 5 μm. (D) Spine plasticity in the peri-infarct area. Box and whiskers plot showing the surviving fraction (SF) ± SEM (*left;* SF_CTRL_= 0.93 ± 0.03%; SF_stroke_= 0.80 ± 0.04%; SF_REHAB_= 0.92 ± 0.02%; one-way ANOVA with post hoc Bonferroni test, *P _CTRL/STROKE_* = 0.003, *P _REAHB/STROKE_* = 0.005), the turnover ratio (TOR) ± SEM (*middle*; TOR_CTRL_= 0.11 ± 0.02%; TOR_STROKE_ = 0.20 ± 0.04%; TOR_REHAB_= 0.10 ± 0.01%; one-way ANOVA with post hoc Bonferroni test, *P* _CTRL/STROKE_ = 0.007, *P _reahb/stroke_* = 0.01) and spine density (SD) ± SEM (*right panel*; SD_CTRL_= 0.39 ± 0.01 μm^−1^; SD*_STROKE_*= 0.24 ± 0.03 μm^−1^; SD_REHAB_= 0.40 ± 0.04 μm^−1^; one-way ANOVA with post hoc Fisher test, *P _CTRL/STROKE_* = 0.006, *P _REAHB/STROKE_* = 0.003) in the peri-infarct area.

As ischemic damage-induced alterations in neural microcircuitry are known to affect dendrites as well as synaptic contacts, producing a large increase in spine turnover in the peri-infarct area(C. E. Brown et al., 2007; Mostany et al., 2010), we next asked whether rehabilitation stabilized synaptic turnover. We thus monitored the appearance and disappearance of apical dendritic spines (Fig. 4C) by performing a frame-by-frame comparison of spines in the mosaic we acquired along the rostro-caudal axis with TPF microscopy. Spines in the peri-infarct region of REHAB animals exhibited increased synaptic stability (Fig. 4D, left panel) and a lower turnover (Fig. 4D, middle panel) than STROKE mice, thereby recapturing features of pre-stroke conditions. The combined rehabilitative treatment also resulted in higher synaptic densities that were comparable to healthy CTRL values (Fig. 4D, right panel). The stabilization of synaptic contacts induced by the rehabilitation intervention was stronger at the proximal level and weakened with increasing distance from the ischemic core (Supplementary Fig. 5D, upper and middle panels).On the other hand, the change in density of spines extended beyond the peri-infarct region (Supplementary Fig. 5D, lower panel).

In brief, the longitudinal imaging of cortical neurons revealed that the combined rehabilitative therapy helped preserving the organization of dendritic arbors in peri-infarct cortex and restored dendritic spine plasticity to control levels.

## Discussion

In the present study, we utilized optical imaging and techniques to manipulate neural activity to obtain a 'big picture' of rehabilitation-induced cortical remodeling in the lesioned and healthy hemispheres of mice with photothrombotic stroke, and then explored synaptic characteristics of these cortical areas. We show that rehabilitative treatment combining physical training and pharmacological inhibition of the contralesional hemisphere stabilized perilesional synaptic contacts, while restoring features of cortical activation typical of pre-stroke conditions, and promoted the formation of an enriched vascular environment to feed the neuronal population that has been newly recruited to motor functions in the peri-lesional area. To the best of our knowledge, our results provide the first evidence of the correlation between rehabilitation-induced neuronal and vascular reshaping in the mouse cortex after stroke.

The rehabilitation paradigm used in this study takes advantage of a combination of robotic rehabilitation and transient silencing of the healthy homotopic hemisphere with BoNT/E. A recent study (Spalletti et al., under revision) tested how synaptic block in the healthy side by BoNT/E could induce improvements in motor functionality. The authors showed that BoNT/E silencing was active in the motor cortex for about 10 days after injection and produced a small and transient improvement in general forelimb motor tests (Gridwalk and Schallert Cylinder test). They hypothesize that silencing of the contralesional motor area stimulates plastic reorganization in perilesional tissue, but the enhancement of plasticity without the guide of an appropriate motor rehabilitation regime is not sufficient to achieve a complete recovery.

Robotic-assisted rehabilitation has received increasing attention in clinical practice in the last years since it allows repeatable and customizable motor training, while ensuring objective measures of motor performance (Balasubramanian, Klein, & Burdet, 2010; Caleo, 2015; Lo, Stephenson, & Lockwood, 2017; Veerbeek, Langbroek-Amersfoort, Wegen, Meskers, & Kwakkel, 2017). Here we employed the M-Platform, a mechatronic device for mouse forelimb training (C. Spalletti et al., 2014) that mimics a robot for upper limb rehabilitation in humans, the “Arm-Guide” (Reinkensmeyer, Takahashi, Timoszyk, Reinkensmeyer, & Kahn, 2001). The same study from Spalletti and colleagues (under revision) demonstrated that this robotic rehabilitation paradigm *per se* was not able to generalize recovery to untrained motor tasks (i.e. Schallert Cylinder test). In our study, robotic rehabilitation alone was not sufficient to restore basic features of cortical activation like amplitude and promptness of the response. Given the uncomplete rescue produced by the single treatment demonstrated both at the behavioral and cortical level, in the following steps of the present study, we focused on the combinatorial therapy that couples BoNT/E treatment with robotic training.

We first investigated the changes induced by the combined rehabilitation paradigm on cortical activity. It has been reported that motor-targeted focal stroke induces abnormally scattered or diffuse structures on cortical motor maps that persist for more than one month after stroke (see, for instance, (Harrison et al., 2013)). The diffuse structure of the motor maps we also observed in non-treated, post-acute stroke afflicted animals (STROKE group) is thought to represent the ongoing reorganization of the underlying cortical circuitry after stroke. In our combined rehabilitation paradigm, the re-establishment of a cluster of neuronal activity in peri-infarct areas reinstated location, timing and amplitude parameters that highly resembled those of healthy control animals. One limitation of the present study is the low spatial resolution of wide-field calcium imaging, where the fluorescence recorded in each pixel is a convolution of dendritic and axonal activation signals from different cortical layers. The high resolution of TPF microscopy might in the future show the origin of the new rehabilitation-induced functional patterns at the single cell level. This technique will allow, for example, determining if the increase in the amplitude of calcium signals we show in REHAB mice is due to a rise in the average activity of few excitatory neurons or if more cells are recruited in the peri-infarct area to control forelimb movements.

The plastic reshaping of cortical motor representations that takes place in the subacute phase after stroke may be linked to the spared or newly sprouted cortico-spinal fibers (Carter et al., 2012; van Meer et al., 2012) or the myelination of transcallosal fibers (van Meer et al., 2012). Large scale imaging techniques like light sheet microscopy (Keller & Dodt, 2012; L Silvestri, Bria, Sacconi, Iannello, & Pavone, 2012) or diffusion magnetic resonance imaging (Chang et al., 2017) of axonal projections directed to and emerging from perilesional neurons will permit answering many questions related to the structural underpinning of functional rewiring.

By performing optogenetic stimulation on the same group of stroked-afflicted, rehabilitated mice, we demonstrated that the progressive spatial and temporal refocusing of motor control in the ipsilesional cortex is associated with increased interhemispheric coupling. The connectivity between homotopic motor cortices was restored 4 weeks after injury in STROKE mice, suggesting that some form of spontaneous recovery compensated for the transcallosal projections lost after stroke. The interhemispheric coupling is further enhanced after the rehabilitative therapy, suggesting a possible correlation between the recovery of the neuronal activation profiles in the perilesional area and the reinforcement of the connectivity with pyramidal neurons that project from the healthy M1. While there are no studies examining, on the same animals, cortical functionality and transcallosal connectivity to draw direct comparisons with, our results are consistent with the hypothesis that rehabilitative training of the paretic upper limb can, on one side, increase the functional activation of motor regions of the ipsilesional cortex (Hodics, Cohen, & Cramer, 2006; Hubbard et al., 2015) and, on the other, decrease interhemispheric inhibition from the contralesional motor cortex (Harris-Love, Morton, Perez, & Cohen, 2011). We speculate that refocusing of the motor representation could be supported by dendritic rewiring in the peri-lesional area (as shown by our *in vivo* imaging result on structural plasticity), as well as by the activity of contralesional pyramidal cells projecting to the peri-infarct area. Further analysis on targeted neuronal populations is needed to clarify how rehabilitation-induced changes in transcallosal response are specifically linked to alterations in the excitatory/inhibitory balance in the ipsilesional spared tissue.

The mouse model of stroke used in this study provides several advantages over more traditional models. Previously, voltage-sensitive dyes has been combined with optogenetics to investigate cortical functional connectivity (Ayling et al., 2009). Recently, this approach applied to stroke models has demonstrated a spontaneous partial recovery of cortical functional connectivity 8 weeks after injury (Lim et al., 2014). Our model, which combines transgenic GCaMP6 mice with AAV-induced expression of ChR2 in the homotopic M1, has the advantage of enabling, on one side, the reliable control and, on the other, the stable monitoring of neuronal activity over weeks and months. We anticipate that the all-optical approach we used will be further extended to the longitudinal exploration of the modified transcallosal axonal projections from the injured to the heathy hemisphere in our rehabilitation paradigm. Moreover, future investigations will help determine how the plasticity of subcortical regions (e.g. basal ganglia, thalamus) is involved in the modulation of rehabilitation-induced motor recovery. Cannula windows (Howe & Dombeck, 2016), fiber-optic systems (Miyamoto & Murayama, 2016; Pisanello et al., 2017) or three photon imaging (Ouzounov et al., 2017) will be optimal tools to image and manipulate neuronal activity in deep structures of the brain.

We hypothesized that the cortical motor circuit reshaping and refinement we observed in rehabilitated mice could be supported by angiogenesis and vascularization. This assumption is based on the relationship existing between the proangiogenic state and neurological improvement in patients with stroke (Arkuszewski, Świat, & Opala, 2009; Hermann & Chopp, 2012; Xiong, Mahmood, & Chopp, 2010). The reason behind it could be that the region of active neovascularization acts as a niche for enhanced neuronal remodeling and tissue repair (Prakash & Carmichael, 2015; Shen et al., 2008). Here, we measured an increase in vascular density after stroke localized in the peri-infarct area that is comparable to healthy animals in stroke non-treated mice (STROKE), while exceed physiological values in mice under double treatment (REHAB). Our results suggest that the transient proangiogenic state induced in response to an ischemic insult is enhanced by rehabilitation. In conjunction with the more efficient neuronal activation we measured in the same areas, our study thus supports the hypothesis that enduring recovery from stroke might result from the association of angiogenesis with neuronal plasticity (Ergul, Alhusban, & Fagan, 2012). We aim to extend the study on vasculature remapping by taking advantage of macro-scale imaging techniques like light-sheet microscopy. This technique, combined with appropriate clearing agents will allow investigating modification of the cytoarchitecture in the entire mouse brain with subcellular resolution (Lugo-Hernandez et al.; L. Silvestri, Mascaro, Costantini, Sacconi, & Pavone, 2014a, 2014b) induced by rehabilitation after stroke.

A commonly agreed hypothesis is that the genesis of an enriched vascular milieu around the infarct area might promote synaptogenesis and dendritic remodeling, in addition to axonal sprouting (Brown et al., 2010; Carmichael, 2006; Kelley & Steward, 1997). In parallel, a recent review by Wahl and colleagues suggested that rehabilitative training might shape the spared and the new circuits by selecting and stabilizing the active contacts and pruning the non-functional ones (Wahl et al., 2014). Alongside with these hypotheses, we show that increased vascular density is associated with increased density of synaptic contacts and stabilization of synaptic turnover induced by the combined rehabilitative treatment. Although there are no other *in vivo* studies examining rehabilitation-induced spine plasticity after stroke to make a comparison with, our results are also in agreement with recent postmortem histological studies where rehabilitation was shown to determine significant increases in spine density of distal apical dendrites in corticospinal neurons (L. Wang, Conner, Nagahara, & Tuszynski, 2016).

Finally, our results show that the combined treatment preserves the random-oriented dendritic architecture at the end of the motor training period. One caveat that our study did not address is whether the random orientation of dendrites and axons was present from an earlier period after stroke, meaning that a shorter training might be sufficient to recover this structural feature. Alternatively, the pre-stroke organization could be recovered after a transient epoch of neurite alignment towards the stroke core. Further investigations will be necessary to resolve these questions.

In this study, we have shown for the first time that combined rehabilitation promotes the recovery of structural features and characteristics of healthy neuronal networks. The restorative therapy rescued the random orientation of pyramidal neurons and supported the stabilization of synaptic contacts. Indeed, rehabilitation counterbalanced the increased synaptic turnover induced by stroke by reinstating baseline dynamics.

Our multi-scale investigation brought to light complementary aspects of the structural and functional plasticity induced by rehabilitation that may lead to the development of more efficient therapies and improve post-stroke recovery in patient populations.

## Materials and methods

### Mice

All procedures involving mice were performed in accordance with the rules of the Italian Minister of Health. Mice were housed in clear plastic cages under a 12 h light/dark cycle and were given *ad libitum* access to water and food. We used two different mouse lines from Jackson Laboratories (Bar Harbor, Maine USA): Tg(Thy1-EGFP)MJrs/J (referred to as GFPM mice) for two-photon imaging experiments and C57BL/6J-Tg(Thy1GCaMP6f)GP5.17Dkim/J (referred to as GCaMP6f mice) for wide-field and optogenetics. Both lines express a genetically-encoded fluorescent indicator controlled by the Thy1 promoter. A subset of GFPM mice imaged for the structural plasticity experiment (dendrites and spines analysis) were used for blood vessels evaluation; a subset of GCaMP6f mice previously used for calcium imaging were analyzed for inter-hemispheric connectivity. Each group contained comparable numbers of male and female mice, and the age of mice was consistent between the groups (4-12 months).

### Photothrombotic stroke induction

All surgical procedures were performed under Zoletil (50 mg/kg) and xylazine (9 mg/kg) anesthesia, unless otherwise stated. After checking by toe pinching that a deep level of sedation had been reached, the animals were placed into a stereotaxic apparatus (Stoelting, Wheat Lane, Wood Dale, IL 60191). The skin over the skull was cut and the periosteum was removed with a blade. The primary motor cortex (M1) was identified (stereotaxic coordinates +1,75 lateral, −0.5 rostral from bregma). Five minutes after intraperitoneal injection of Rose Bengal (0.2 ml, 10 mg/ml solution in Phosphate Buffer Saline (PBS); Sigma Aldrich, St. Louis, Missouri, USA), white light from an LED lamp (CL 6000 LED, Carl Zeiss Microscopy, Oberkochen, Germany) was focused with a 20X objective (EC Plan Neofluar NA 0.5, Carl Zeiss Microscopy, Oberkochen, Germany) and used to illuminate the M1 for 15 min to induce unilateral stroke in the right hemisphere. Afterwards, the skin over the skull was sutured and the animals were placed in recovery cages until full recovery. The CTRL animals are not subjected to photothrombosis.

### Optical windows

For the experiments on GCaMP6f mice, we performed a thinned skull preparation on the right hemisphere between bregma and lambda to create an optical window. After applying the local anesthetic lidocaine 2% (20 mg/mL), the skin over the skull and periosteum was removed. The skull over most of the right hemisphere was thinned using a dental drill. A cover glass and an aluminum head-post were attached to the skull using transparent dental cement (Super Bond, C&S). We waited at least 4-5 days after the surgery for the mice to recover before the first imaging session.

For the experiments on GFPM mice, we created a square (3×5 mm^2^) cranial window centered laterally on the right M1 (+1.75 mm from bregma) and extending rostro-caudally from 1 mm posterior to the bregma to lambda. The protocol we followed for cranial window preparation was slightly modified from Holtmaat et al., 2009(Holtmaat et al., 2009) and Allegra Mascaro et al., 2014(Allegra Mascaro, Sacconi, & Pavone, 2014). Briefly, we administered anesthetized mice a subcutaneous injection of dexamethasone (0.04 ml per 2 mg/ml). The animals were then placed into a stereotaxic apparatus; after applying the local anesthetic lidocaine 2% (20 mg/mL), the skin over the skull was removed. Using a dental drill (Silfradent, ForlÌ-Cesena Italia), the border of the area of interest was thinned and the central part of the bone was then gently removed. The exposed brain was covered with a circular cover glass; the optical window was sealed to the skull with a mixture of dental cement and acrylic glue. Finally, an aluminum head-post was attached onto the skull using dental cement (Super Bond, C&S, Sun medical Moriyama City, Shiga, Japan). The surgery was followed by the first imaging session under the two-photon microscope. If the cranial windows were opaque on the second imaging session, the windows were removed and cleaned, and imaging was performed immediately afterwards. After the last imaging session, all animals were perfused with 150 mL of Paraformaldehyde4% (PFA, Aldrich, St. Louis, Missouri, USA).

### Intracortical injections

We used a dental drill to create a small craniotomy over M1, which was identified by stereotaxic coordinates. Botulinum Neurotoxin E (BoNT/E) injections were performed during the same surgical session in which the photothrombotic lesions were created. We injected 500 nl of BoNT/E (80 nM) divided in 2 separate injections of 250 nl at (i) +0.5 anteroposterior, +1.75 mediolateral and (ii) +0.4 anteroposterior, +1.75 mediolateral at 700 μm cortical depth. For virus injections, we delivered 1 μl of AAV9-CaMKII-ChR2-mCherry (2.48*10^13^ GC/mL) 700-900 μτ deep inside the cortex. The skin over the skull was then sutured; the animals were placed in a heated cage (temperature 38°) until they fully recovered.

### Motor training protocol on the M-Platform

Mice were allowed to become accustomed to the apparatus before the first imaging session so that they became acquainted with the new environment. The animals were trained by means of the M-Platform, which is a robotic system that allows mice to perform a retraction movement of their left forelimb(C. Spalletti et al., 2014). Briefly, the M-Platform is composed of a linear actuator, a 6-axis load cell, a precision linear slide with an adjustable friction system and a custom-designed handle that is fastened to the left wrist of the mouse. The handle is screwed onto the load cell, which permits a complete transfer of the forces applied by the animal to the sensor during the training session. Each training session was divided into “trials” that were repeated sequentially and consisted of 5 consecutive steps. First, the linear actuator moved the handle forward and extended the mouse left forelimb by 10 mm (full upper extremity extension). Next, the actuator quickly decoupled from the slide and a tone lasting 0.5 s informed the mouse that it should initiate the task. If the animal was able to overcome the static friction (approximately 0.2 N), it voluntarily pulled the handle back by retracting its forelimb (i.e. forelimb flexion back to the starting position). Upon successful completion of the task, a second tone that lasted 1 sec was emitted and the animal was given access to a liquid reward, i.e. 10 μl; of sweetened condensed milk, before starting a new cycle.

To detect the movement of the wrist of the animal in the low-light condition of the experiment, an infrared (IR) emitter was placed on the linear slide, and rigidly connected to the load cell and thus to the animal's wrist. Slide displacement was recorded by an IR camera (EXIS WEBCAM #17003, Trust) that was placed perpendicular to the antero-posterior axis of the movement. Position and speed signals were subsequently extracted from the video recordings and synchronized with the force signals recorded by the load cell (sampling frequency = 100 Hz).

All groups performed at least one week (5 sessions) of daily training, starting 26 days after injury for STROKE mice, 5 days after stroke for the ROBOT and REHAB group and after the surgery for CTRL animal.

### Behavioral evaluation of forelimb use asymmetry

Spontaneous forelimb use was assessed in mice via the Schallert cylinder test. This test has been developed in rats and adapted for mice(Baskin, Dietrich, & Green, 2003; Schallert et al., 2000). Briefly, mice were placed individually inside a clean cylinder (diameter, 8 cm; height, 20 cm) that was placed vertically above a transparent surface and were allowed to explore this enclosure for 5 minutes. Mice were video-taped with a camcorder (C270, Logitech) that was positioned underneath the transparent surface. The number of paw touches on the wall during rearing was recorded offline, and the percentage use of each paw relative to the total amount of touches was calculated.

### Optogenetic stimulation and simultaneous recording of GCaMP6f activity

After the last training session (i.e. 30 days after stroke and at least 2 weeks after the AAV injection in the CTRL group) mice were anesthetized under Zoletil (50 mg/kg) and xylazine (9 mg/kg) and placed into the stereotaxic holder. A small (2×2 mm^2^) craniotomy was performed over the injected area. After placing the mouse under the wide field fluorescence microscope, we performed repeated laser (473 nm) stimulation (1-2 Hz, pulse duration 3-5 ms, pulse train duration 5 sec, laser power at the focal plane 5 mW) on the left M1, which was localized by mCherry fluorescence. Spurious activation of ChR2 from the green LED (used for GCaMP6f fluorescence excitation) was avoided by blocking half the illumination path with a shutter positioned after the collimator.

### Wide-field fluorescence microscopy

The custom-made wide-field imaging setup was equipped with two excitation sources for the simultaneous imaging of GcaMP6f fluorescence and light-stimulation of ChR2. For imaging of GcaMP6f fluorescence, a 505 nm LED (M505L3 Thorlabs, New Jersey, United States) light was deflected by a dichroic filter (DC FF 495-DI02 Semrock, Rochester, New York USA) on the objective (2.5× EC Plan Neofluar, NA 0.085, Carl Zeiss Microscopy, Oberkochen, Germany). A 3D motorized platform (M-229 for xy plane, M-126 for z-axis movement; Physik Instrumente, Karlsruhe, Germany) allowed sample displacement. The fluorescence signal was selected by a band pass filter (525/50 Semrock, Rochester, New York USA) and collected on the sensor of a high-speed complementary metal-oxide semiconductor (CMOS) camera (Orca Flash 4.0 Hamamatsu Photonics, NJ, USA).

To perform optogenetic stimulation of ChR2, a 473 nm continuous wavelength (CW) laser (OBIS 473nm LX 75mW, Coherent, Santa Clara, California, United States) was overlaid on the imaging path using a second dichroic beam splitter (FF484-Fdi01-25x36, Semrock, Rochester, New York USA). The system has a random-access scanning head with two orthogonally-mounted acousto-optical deflectors (DTSXY400, AA Opto-Electronic, Orsay France). A 20X objective (LD Plan Neofluar, 20×/0.4 M27, Carl Zeiss Microscopy, Oberkochen, Germany) was used to demagnify the image onto a 100X100 pxl^2^ area of the sCMOS camera sensor (OrcaFLASH 4.0, Hamamatsu Photonics, NJ, USA). Images (512×512 pixels, pixel size 9 μm) were acquired at 25 Hz.

### Two-photon fluorescence microscopy

The custom made apparatus for two-photon microscopy included a mode-locked Ti: Sapphire laser (Chameleon, Coherent Inc.) that supplied the excitation light. The laser beam was scanned in the xy-plane by a galvo system (VM500, GSI Lumonics). An objective lens (XLUM 20X, NA 0.95, WD 2 mm, Olympus) focused the beam onto the specimen. A closed-loop piezoelectric stage (PIFOC ND72Z2LAQ, PhysikInstrumente, Karlsruhe Germany) allowed axial displacements of the objective up to 2 mm with micrometric precision. Finally, the fluorescence signal was collected by a photomultiplier tube (H7710-13, Hamamatsu Photonics). Custom-made software was developed in LabVIEW 2013 (National Instruments).

### Labelling of brain vasculature

The vasculature was stained using the protocol described by Tsai et al.(Tsai et al., 2009), except that we replaced fluorescein (FITC)-conjugated albumin with 0.05% (w/v) tetramethylrhodamine (TRITC)-conjugated albumin (A23016, Thermo Fisher Scientific, Waltham, Massachusetts, USA) in order to avoid spectral overlap between GFP and FITC. Under deep anesthesia, mice were transcardially perfused first with 20-30 ml of 0.01 M PBS (pH 7.6) and then with 60 ml of 4% (w/v) paraformaldehyde (PFA) in PBS. This was followed by perfusion with 10 ml of fluorescent gel. After perfusion, a low temperature (ice cold) was maintained to ensure rapid solidification of the gel. After 30 min of cooling, the brain was carefully extracted to avoid damage to pial vessels, and it was incubated overnight in 4% PFA at 4 °C. Dissected cortices were cleared with thiodiethanol (166782 TDE, Sigma Aldrich, St. Louis, Missouri, USA)(Aoyagi, Kawakami, Osanai, Hibi, & Nemoto, 2015; Costantini et al., 2015; Staudt, Lang, Medda, Engelhardt, & Hell, 2007), specifically with serial incubation in 30% and 63% TDE/PBS for 1 h and 3 h, respectively, at room temperature. The cleared cortices were flattened using a quartz coverslip #No1 (UQG Optics).

### Image analysis

#### Wide-field calcium imaging

During each experimental session, the mouse's head was restrained and placed on the M-platform under the wide-field microscope. To avoid head movement artefacts, each frame of the fluorescence stack was offline registered by using two reference points (corresponding to bregma and lambda) that were previously marked on the glass window during the surgery procedure.

For each stack (*FluoSt*) a median time series of GCamp6f fluorescence signal (*mF*) was extracted, where the value of mF at each *i*-*th* time point corresponded to the median value computed on all the pixels of the *i*-*th* frame of *FluoSt*. The *mF* was then oversampled and synchronized to the 100 Hz force and position signals.

The *mF* was used to define a GCamp6f fluorescence signal baseline ***F***_**0**_, which was identified by the concomitant absence of fluorescence and force signal deflections. ***F***_**0**_ was selected within a 2.5 ± 0.7 s interval (***I***) of 62 frames where the fluorescence signal was below 1 standard deviation (STD) of the whole *mF* signal and the corresponding force signal showed a value below of 1 STD of the whole recorded force signal. The fluorescence signal interval was used to reconstruct a 512×512 matrix, i.e. baseline matrix, in which the value of each pixel *B* of the {*m*,*n*} coordinates of the matrix was computed as follows:

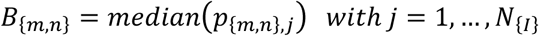

where ***P*****{*_**m,n**_*}**,***j*** is the value of the pixel *p* of the {*m*,*n*} coordinates at the *j*-*th* frame of the interval ***I*** and ***N***_**{*****I*****}**_ is the length of Interval ***I***. The baseline matrix was then used to normalize all the frames of the fluorescence stack *FluoSt*.

The noise-threshold of 1 STD of the whole recorded force signal was used as a measure of the force peaks exerted by the animal during the retraction task. According to Spalletti et al. 2013(M. Spalletti et al., 2013), a force peak is defined as force values that transiently exceed the noise-threshold and result in a movement of the linear slide, as detected by variation of the position signal. To maintain consistency in this analysis, peaks of force that did not result in a movement of the slide were not considered.

The onset of each force peak was used as reference time point to select a sequence of 60 frames (2.4 s, where 0.4 s preceded the force peak) from the *FluoSt*. All sequences were visually checked to exclude possible spurious activation (e.g. early activation or no activation) from the analysis. All the selected sequences (*Seqs*) of the animal *An* on day *d* were compiled, defining a *stack of Seqs,* to compute the *Summed Intensity Projection* for the *An* at *d* (*SIP_An d_*). The SIP is a matrix of 512x512 pixels, in which the value *P* of the pixel of the {*m*, *n*} coordinates is computed as follows:

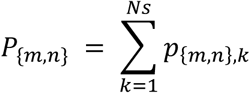

wherewhere ***P***_**{*****m,n*****}**_,***k*** is the value of the pixel p of the {*m*, *n*} coordinates at the *k*-*th* frame of *stack of Seqs*, and ***N_s_*** is the number of frames of the *stack of Seqs*.

The most active area of the *SIP_An d_* was then detected by thresholding the *SIP_An d_* with a *median (SIPAn d)* + *STD (SIPAn d)* threshold value. The threshold *SIP_An d_* (*th*-*SIP_An d_*) computed for each week of training on the M-Platform (d=1,…,4 of week W) was superimposed and the common areas, activated at least for 3 dailysessions out of 5 (60%), were labeled as “regions of interest” (ROIs) of the *SIPAn*. We further refined the analysis by dividing the image into two areas, and identifying one anterior ROI [−0.25 – +1.95 mm from bregma (B), AP] and one posterior ROI[+1.95 – + 4.15 mm from B, AP].

We also used the ROIs defined for each individual animal to identify the average ROIs among mice from the same experimental group: *ROI_g_* with g = “CTRL”, “STROKE”, “ROBOT”, “REHABlw” and “REHAB4w” (see Supplementary Fig. 2B). Thus, the ROIs from animals of the same group were compiled and further thresholded (60%) to define the *ROI_g_*. The extent of the *ROI_g_* was computed as follows:

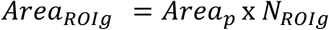

where ***Area**_**p**_* corresponds to the area of a single pixel of the image (0.0086 mm^2^) and ***N**_**R0I_g_**_* is the number of pixels composing the *ROI_g_*. Moreover, the centroid of each *ROI_g_* was identified and its Euclidean distance from bregma was computed (Fig. 2F).

The ROIs defined for each individual animal were further used to extract the GCamp6f fluorescence signal corresponding to the activity of those areas. Indeed, from each frame of the *FluoS*, only pixels belonging to the selected ROI were considered when calculating the representative median value. Thus, a median time series, ***F**_**R0I**_*, was extracted from the whole *FluoS* and was representative of the ROI. The fluorescence signal was normalized 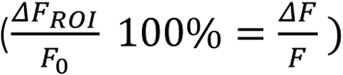 and low-pass filtered to clean the signal from the detected breathing artefacts (Chebyshev filter with cutting frequency = 9 Hz).

The previously detected force peaks were then used to select the GCamp6f fluorescence peaks from the 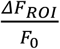 signal. A time window that lasted 4 seconds, i.e. *wnd,* and was centered at the onset of the force peak, was used to delimit a part of the 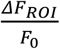 signal, i.e. ***F**_**wnd**_*, to identify the corresponding fluorescence peak. A fluorescence peak was defined as the part of ***F**_**wnd**_* that overcame the value of *median+3STD* calculated for the whole signal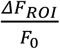. The onset of the peak was detected as

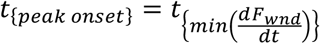

calculated for the [***t_st_ t_max_***] time interval, where ***t_st_*** and ***t_max_*** are the time points corresponding to the start of the *wnd* and the maximum of ***F_wnd_***, respectively.

From the fluorescence and force peaks, different parameters were computed as follows:

- the maximum of the peak (*Peak amplitude*).
- the *Slope* of the peak was defined as follows:

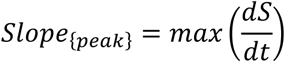

between the *t*_{*s peak onset*_] and *t_S max_* where *S* = *fluo* or *force* signal.

- the time delay between the occurrence of the maximum of the fluorescence (*ΔT*).

The first movement time point was defined as the first variation of the position signal 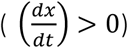 detected along the *wnd* interval.

#### Optogenetically-induced calcium activity

On every stack, we analyzed the calcium activity averaged over a round ROI (2.3 mm of diameter) centred on the peri-infarct area. Changing in the variation of fluorescence signal ΔF/F) triggered by light irradiation of the contralateral hemisphere that were below 1% were excluded from the analysis. A calcium response was considered optogenetically-triggered if it started within 120 ms of optogenetic stimulation. The activation delay refers to the average (± Standard Error of the Mean, SEM) delay of the calcium peak with respect to the onset of laser stimulation. The success rate reports the number of times the laser stimulation successfully triggered contralateral activation over the total number of stimulation trials.

#### Two-photon imaging

For vessel orientation analysis, 1 mm deep stacks with 3 μm z-steps were acquired by TPF microscopy. From these acquisitions throughout the entire cortical depth of the right hemisphere, we extracted maximum intensity projections (MIPs; 300 μm thick) over selected substacks, avoiding meningeal vessels in the superficial 100 μm Four MIPs of proximal regions (within 500 μm from the stroke core), and four MIPs of distal regions (1-1.5 mm from the core) for each mouse were analyzed. Polar plots of vessel orientation were obtained by measuring frame by frame the angle between each structure and the rostro-caudal axis (n = 30 vessels for each group). For the vessel density analysis, the stacks were first binarized using automatic thresholding with ImageJ. The sum of the pixel count from histograms of the binarized stacks was used as a measure of blood vessel density.

For the structural plasticity analysis of dendrites and spines of pyramidal neurons, we compared stacks in a vertical mosaic acquired in the rostro-caudal direction. During the last week of training (1st and 4th day), we acquired a mosaic of 100 μm thick stacks (113 × 113 μm^2^) distanced 200 μm from each other along the rostro-caudal axis starting from the borders of the stroke core. We grouped stacks near (<500 μm) and far (from 1000 to 1500 μm from the core, namely proximal and distal regions, respectively. Dendrite orientation was evaluated on 15 dendrites for each stack (on average) by measuring frame by frame the angle between each structure and the rostro-caudal axis. The stroke core was considered to be at 0°. For synaptic plasticity analysis, the fluorescence signal of a spine had to be be at least 1 standard deviation higher than the dendritic shaft fluorescence to be included in the analysis. We quantified the plasticity of dendritic spines using two functions: surviving fraction (SF) and turnover ratio (TOR) (Holtmaat et al., 2009); the SF describes the fraction of persistent structures:

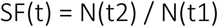

where N(t1) is the number of spines present during the first imaging session (26 days after injury for STROKE and REHAB mice), while N(t2) indicates those structures that present 4 days after the first imaging session. The TOR evaluates the fraction of newly appeared in the images and disappeared structures:

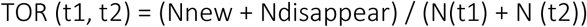

where Nnew is the number of structures that are reported for the first time at time t2, Ndisappear is the number of structures that were present at time t1 but which are no longer present at time t2; N (t1) and N (t2) are the total of spines present on t1 and t2, respectively. Unless otherwise stated, data are reported as mean ± SEM.

## Acknowledgement

We thank Alessio Masi and Marie Caroline Muellenbroich for very useful discussion about the manuscript; Giuseppe De Vito for assistance on statistics analysis. This project has received “funding from the H2020 EXCELLENT SCIENCE - European Research Council (ERC) under grant agreement ID n. 692943 BrainBIT”. In addition, it was supported by the European Union′s Horizon 2020 research and innovation program under grant agreements No. 720270 (Human Brain Project) and 654148 (Laserlab-Europe). The project was also supported by the Italian Ministry for Education, University, and Research within the framework of the Flagship Project NanoMAX, and by “Ente Cassa di Risparmio di Firenze” (private foundation). Part of this work was performed within the framework of the Proof of Concept Studies for the ESFRI research infrastructure project Euro-BioImaging at the PCS facility LENS.

---

**Supplementary Figure 1.**
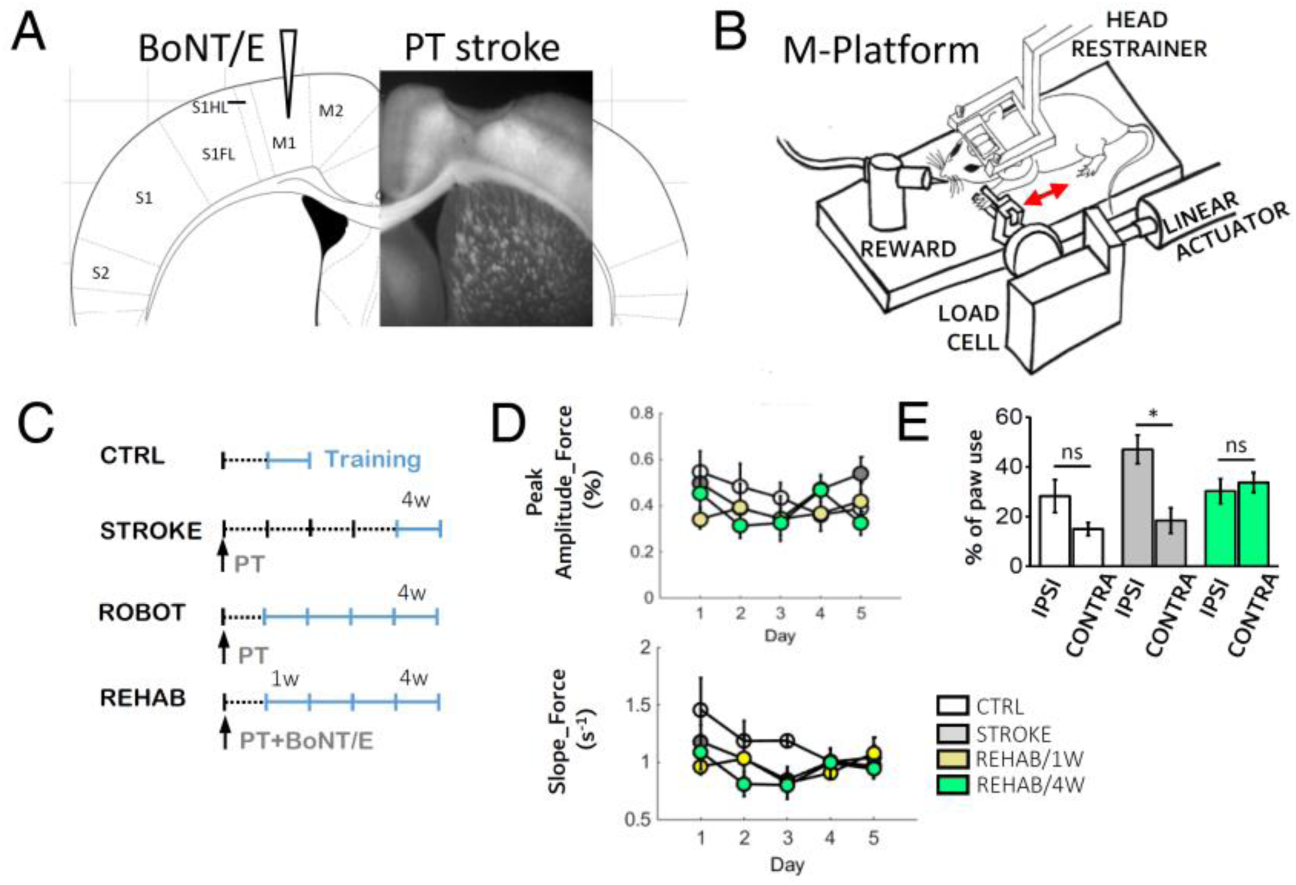
Experimental design. (A) A schematic of the experimental protocol, which combines the photothrombotic stroke in the primary motor cortex (M1) with a contralesional injection of BoNT/E into the homotopic cortex. (B) A schematic representation of the M-Platform that was used for rehabilitative training. (C) The experimental timeline for the CTRL, STROKE and REHAB groups in the awake imaging experiment. Light blue lines refer to training weeks. W = week; PT = photothrombosis. (D) Graphs showing the Peak amplitude (left) and Slope (right) of the force signals recorded in the pulling phase during training on the M-Platform over 5 days (4 weeks after injury for STROKE mice, 1 and 4 weeks after injury for REHAB mice, during the week of training for CTRL mice). There is no significant difference between the groups and over the 5 days. (E) Histogram showing the % of ipsilateral versus contralateral paw use in the Schallert cylinder test performed 30 days post stroke by GCaMP6f mice (Nmice_CTRL_= 3, Nmice_STROKE_ = 3, Nmice_REHAB_= 3, CTRL_IPSI_=28 ± 7%; CTRL_CONTRA_=15 ± 3%; STROKE_IPSI_=47 ± 6%; STROKE_CONTRA_=18 ± 5%; REHAB_IPSI_= 30 ± 5%; REHAB_CONTRA_ = 34 ± 4%; t-test STROKE _IPSI_/STROKE_CONTRA_* *P* = 0.01). IPSI and CONTRA refer to the ipsilesional and contralesional paws, respectively; ns, not significant.

**Supplementary Figure 2.**
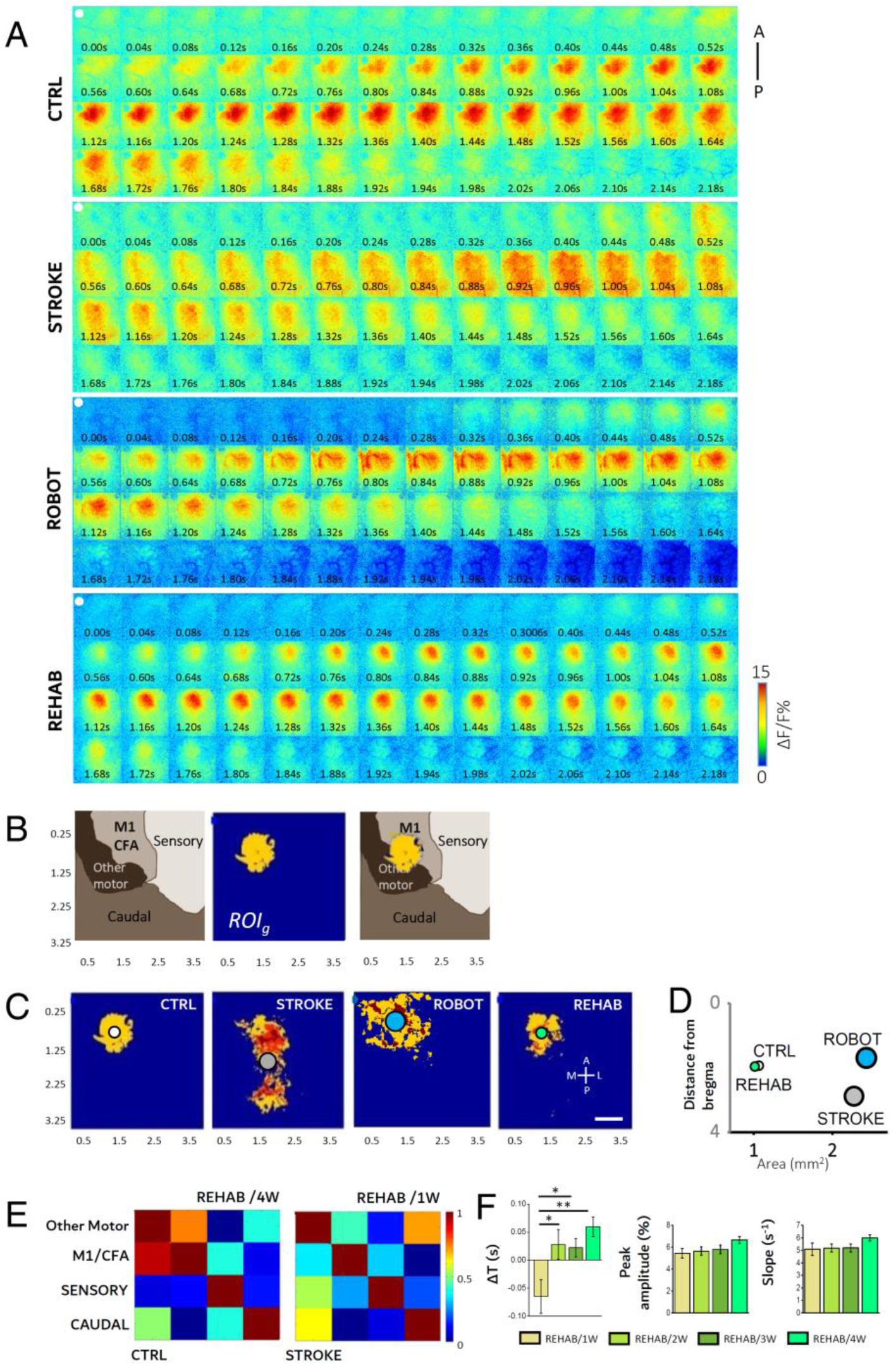
Wide-field imaging of cortical activation profiles over 4 weeks of rehabilitation. (A) Complete image sequences of cortical activation as assessed by calcium imaging during pulling of the handle by the contralateral forelimb of CTRL (top), STROKE (middle) and REHAB (bottom) GCaMP6f mice in the M-Platform. A-P, anterior-posterior. The white dot indicates bregma. Scale bar = 1 mm. (B) The image on the left shows a functional map based on the intracortical microstimulation (ICMS) studies of Tennant (2011, Cerebral Cortex) and Alia (2016, Sci Reports) that was used as a reference map. The middle panel shows the average thresholded ROI (*ROI_g_*) computed for CTRL animals during voluntary contralateral forelimb pulling in the M-Platform. The image on the right shows a merged image of the functional map and *ROIg*. CFA,caudal forelimb area. The caudal area includes visual and associative regions. (C) The panels show the average thresholded ROI (*ROI_g_*) computed for each experimental group. The circles represent the centroids of the *ROI_g_* for CTRL (white), STROKE (light gray), ROBOT (blue) and REHAB (green) groups. Scale bar = 1 mm. (D) The areas of the *ROI_g_* and their distance from bregma are presented in the scatter plot. (E) Examples of partial correlation matrices of cortical activation during voluntary pulling. The correlation analysis on REHAB mice was performed based on two evaluation time points: one week (REHAB/1W) and four weeks (REHAB/4W) after stroke. (F) Longitudinal analysis of cortical activation profiles over the four rehabilitation weeks. Graphs in the left, middle and right panels are calculated as in Fig. 1E, G, and H, respectively. Left panel: Delays in cortical activation in caudal regions following the forelimb retraction task are reported for the 4 weeks of rehabilitative training (Nmice_REHAB_= 6; ΔT_REHAB/1w_ = −0.06 ± 0.03 s, ΔT_REHAB/2w_ = −0.03 ± 0.03 s, ΔT_REHAB/3w_ = 0.02 ± 0.02 s, ΔT_REHAB/4w_ = 0.06 ± 0.02 s; one-way ANOVA followed by the Tukey test: ** *P* <0.01, *P <0.05; REHAB/4W here corresponds to the REHAB group in all the other panels). Middle panel: the maximum of calcium imaging fluorescence peaks for the 4 weeks of rehabilitative training (Nmice_REHAB_= 6; Peak amplitude_REHAB/1w_= 5.4 ± 0.4%, Peak amplitude_REHAB/2w_ = 5.6 ±0.4 %, Peak amplitude_REHAB/3w_ = 5.8 ± 0.4%, Peak amplitude_REHAB/4w_ = 6.6 ± 0.3 %; No significant changes were observed over the 4 weeks period). Right panel: The graph shows the slope (average ± SEM) of the calcium imaging fluorescence in the rising phase (Nmice_REHAB_= 6; Slope_REHAB/1w_= 5.1 ±0.5 s^−1^, Slope_REHAB/2w_= 5.2 ±0.3 s^−1^, Slope_REHAB/3w_= 5.2 ±0.3 s^−1^, Slope_REHAB/4w_= 6.0 ±0.2 s^−1^; No significant changes were observed over the 4 weeks period).

**Supplementary Figure 3.**
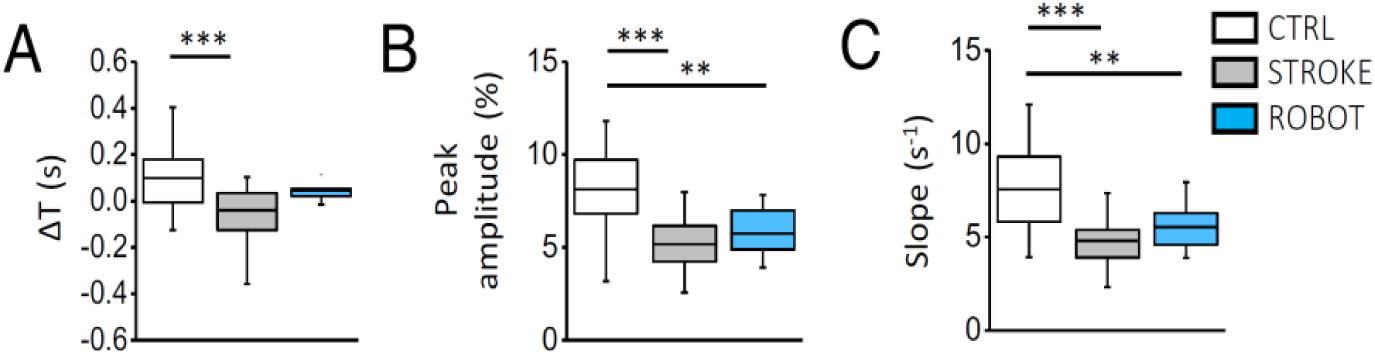
Motor training on the M-platform is not sufficient to trigger recovery of cortical activity patterns. (A) The delays in cortical activation in caudal regions in response to contralateral forelimb retraction are shown for the 3 groups (Nmice_CTRL_= 4, Nmice_STROKE_ = 6, Nmice_ROBOT_= 3; ΔT_CTRL_ = 0.10 ± 0.03 s, ΔT_STROKE_= −0.04 ± 0.02 s, ΔT_ROBOT_= 0.05 ± 0.01 s; one-way analysis of variance (ANOVA) followed by the Bonferroni test: *** *P* = 0.0007). (B) The graph shows the maximum of the fluorescence peaks from the same calcium traces as in (A) (Peak amplitude_CTRL_ = 8.1 ± 0.5 %; Peak amplitude_STROKE_ = 5.2 ± 0.3%; Peak amplitude_ROBOT_ = 5.7 ± 0.5%; one-way ANOVA followed by Facqthe Bonferroni test: *** *P* = 9*10^−7^, ** *P* = 0.004). (C) The graph shows the slope of the fluorescence in the rising phase of the trace (Slope_CTRL_ = 7.5 ± 0.5 s-1; Slope_STROKE_ = 4.8 ± 0.2 s-1, Slope_ROBOT_ = 5.5 ± 0.3 s-1; one-way ANOVA followed by the Bonferroni test: *** *P* =6*10^−7^, ** *P* = 0.02). All values are reported as average ± SEM.

**Supplementary Figure 4.**
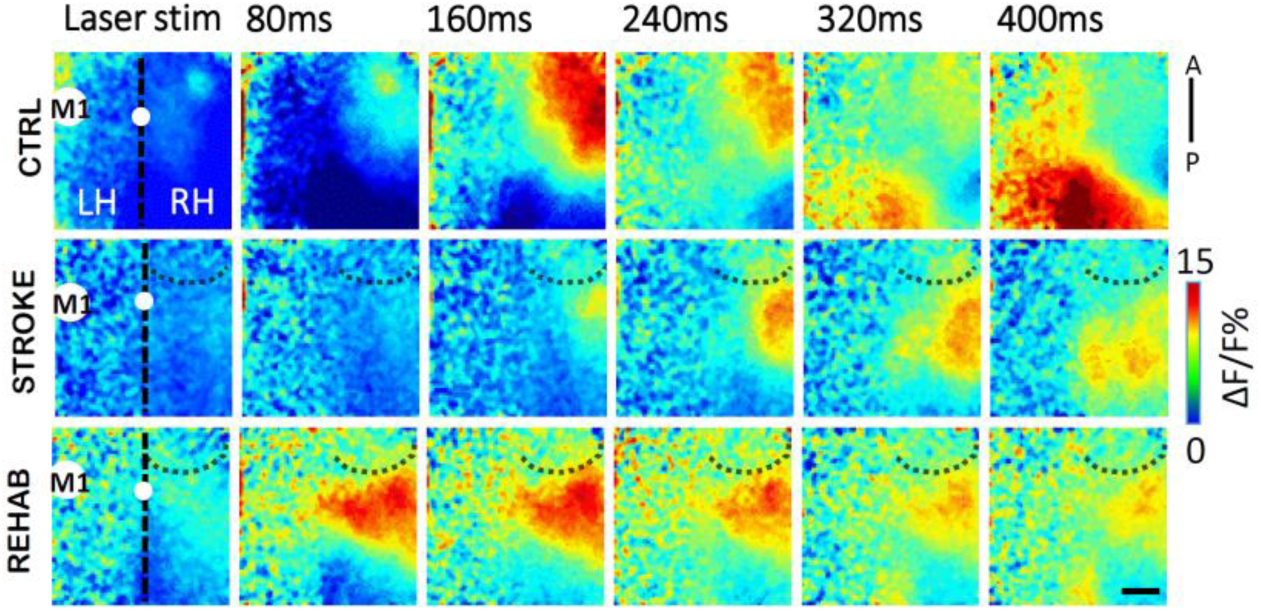
Optogenetic activation of contralateral hemisphere. Representative image sequences from a single animal in each group showing ipsilesional cortical activation as assessed by calcium imaging following 473 nm laser stimulation (1 Hz, 5ms pulse duration) of the ChR2-expressing intact M1. The white spot on the vertical black dotted line (midline) indicates bregma. The dotted curve lines highlight the area injured by stroke. A,P: anterior, posterior. LH, left hemisphere; RH, right hemisphere. Scale bar = 1 mm.

**Supplementary Figure 5.**
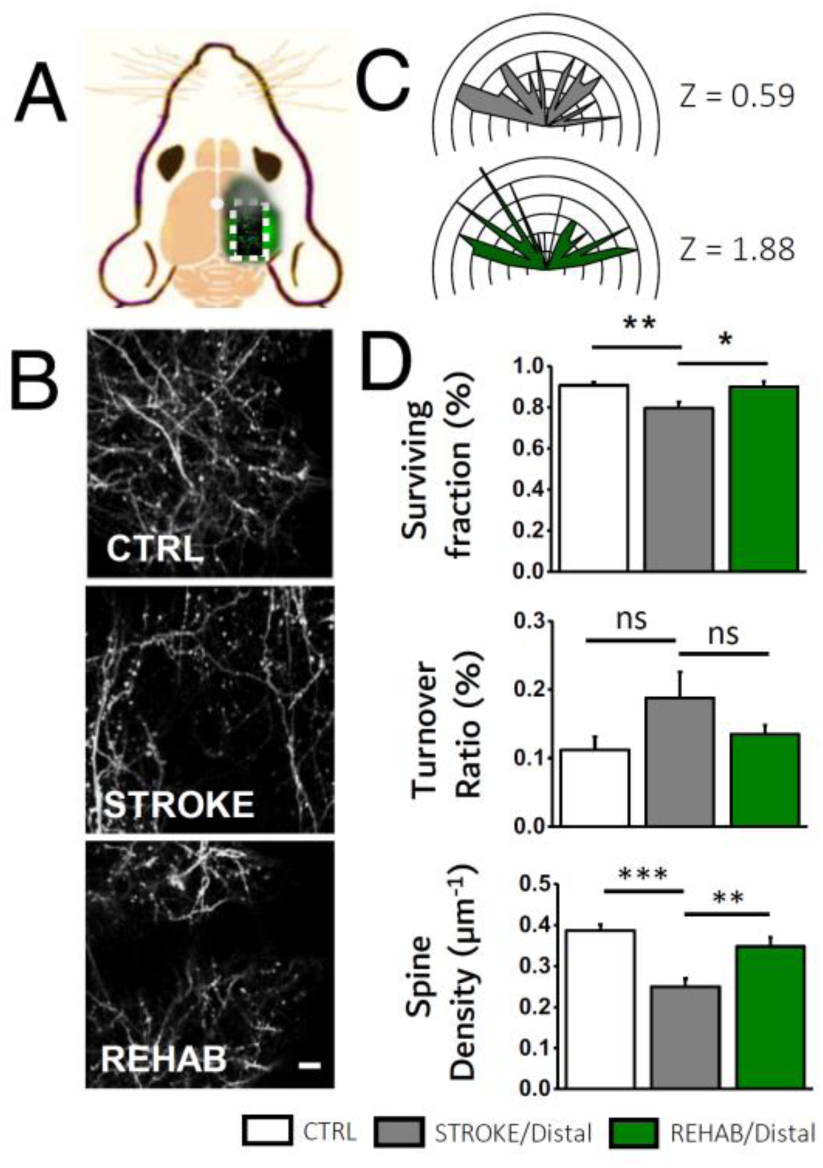
In vivo imaging of dendritic and spine plasticity in regions proximal and distal from stroke core. (A) A schematic representation of the field of view (i.e. area within the white dotted square) of two-photon imaging of dendritic and spine plasticity in GFPM mice. The red spot indicates the site of the stroke lesion. (B) Representative examples of maximum intensity projection of two-photons stacks of dendritic branch orientation in the peri-infarct area in the CTRL, STROKE and REHAB groups. Scale bar = 10 μm. (C) Polar plots showing the angular distribution of dendrites in the distal area (Nmice_CTRL_= 5, Nmice_STROKE_ = 4, Nmice_REHAB_= 3). Z scores, calculated by the Rayleigh Test for circular statistics, are reported for each experimental class on the right of the polar plot; ns, not significant). (D) Histograms showing the SF *(upper panel;* Nmice_CTRL_= 6, Nmice_STROKE_ = 4, Nmice_REHAB_= 3; SF_CTRL_= 0.92 ± 0.02%; SF_STROKE Distal_= 0.80 ± 0.03%; SF_REHAB DIstal_= 0.90 ± 0.03%; one way ANOVA with post hoc Bonferroni: ** *P* =0.007, * *P* =0.023), TOR (*lower panel*; *Nmice_CTRL_*= 6, *Nmice_STROKE_* = 4, *Nmice_REHAB_*= 3; TOR_CTRL_= 0.11 ± 0.02%; TOR_STROKE Distal_ = 0.19 ± 0.04%; TOR_REHAB_= 0.13 ± 0.01% t-test: one way ANOVA with post hoc Bonferroni: not significant for all comparisons) and spine density (SD) (*right panel*; Nmice_CTRL_= 3, Nmice_STROKE_ = 3, Nmice_REHAB_= 3; SD_CTRL_= 0.39 ± 0.01 μm^−1^; SD_STROKE_= 0.25 ± 0.02 μm^−1^; SD_REHAB_= 0.35 ± 0.02 μm^−1^; one-way ANOVA with post hoc Fisher test, *P CTRL/STROKE* = 0.0001, *PCTRL/STROKE* = 0.005) in the distal region (>1000 μm from the stroke core). Data are means ± SEM.

